# Task space exploration improves adaptation after incompatible virtual surgeries

**DOI:** 10.1101/2021.08.06.455370

**Authors:** Denise J. Berger, Daniele Borzelli, Andrea d’Avella

## Abstract

Humans have a remarkable capacity to learn new motor skills, a process that requires novel muscle activity patterns. Muscle synergies may simplify the generation of muscle patterns through the selection of a small number of synergy combinations. Learning new motor skills may then be achieved by acquiring novel muscle synergies. In a previous study, we used myoelectric control to construct virtual surgeries that altered the mapping from muscle activity to cursor movements. After compatible virtual surgeries, which could be compensated by recombining subject-specific muscle synergies, participants adapted quickly. In contrast, after incompatible virtual surgeries, which could not be compensated by recombining existing synergies, participants explored new muscle patterns, but failed to adapt. Here, we tested whether task space exploration can promote learning of novel muscle synergies, required to overcome an incompatible surgery. Participants performed the same reaching task as in our previous study, but with more time to complete each trial, thus allowing for exploration. We found an improvement in trial success after incompatible virtual surgeries. Remarkably, improvements in movement direction accuracy after incompatible surgeries occurred faster for corrective movements than for the initial movement, suggesting that learning of new synergies is more effective when used for feedback control. Moreover, reaction time was significantly higher after incompatible than after compatible virtual surgeries, suggesting an increased use of an explicit adaptive strategy to overcome incompatible surgeries. Taken together, these results indicate that exploration is important for skill learning and suggest that human participants, with sufficient time can learn new muscle synergies.

**NEW & NOTEWORTHY:** Motor skill learning requires the acquisition of novel muscle patterns, a slow adaptive process. Here we show that learning to control a cursor after an incompatible virtual surgery, a complex skill requiring new muscle synergies, is possible when enough time for task space exploration is provided. Our results suggest that learning new synergies is related to the exceptional human capacity to acquire a wide variety of novel motor skills with practice.

## INTRODUCTION

It is everyone’s experience that practice is the key to learning new skills. Learning a new motor skill often requires the generation of novel muscle activity patterns and it associated to plastic changes in the CNS (Dayan and Cohen 2011). Understanding the mechanisms employed by the CNS to control movements, coordinate many degrees-of-freedom, and learn new motor skills represents a fundamental challenge in neuroscience. Muscle synergies have been proposed as modules that simplify the processes of force generation and movement control by reducing the number of parameters that the CNS must regulate when generating muscle patterns (d’Avella et al. 2003; d’Avella and Pai 2010; Bizzi et al. 2002, 2008; Flash and Hochner 2005; Giszter et al. 2007; Jacobs and Macpherson 1996; Lacquaniti et al. 2012; Ting and McKay 2007; Tresch et al. 1999). Initial evidence supporting such a modular architecture for motor control has come mainly from muscle pattern decomposition, i.e., from the observation that by combining a few muscle synergies, muscle patterns can be reliably reconstructed (d’Avella et al. 2003, 2006; d’Avella and Lacquaniti 2013; d’Avella and Pai 2010; Bizzi et al. 2008; Bizzi and Cheung 2013; Cheung et al. 2005; Dominici et al. 2011; Giszter 2015; Ivanenko et al. 2004; Ting and McKay 2007; Torres-Oviedo and Ting 2007; Tresch et al. 2006). However, whether muscle synergies might be due to task and biomechanical constraints is still debated (Kutch and Valero-Cuevas 2012; Tresch and Jarc 2009).

Recently, we obtained direct support for the synergy hypothesis by perturbing the muscle-to-force mapping in a virtual reality setup using myoelectric control (Berger et al. 2013). We created perturbations altering the mapping in a way that could either be compensated by recombining existing subject-specific muscle synergies (compatible perturbations) or could not (incompatible perturbations). We refer to these perturbations as *virtual surgeries*, as they simulate in a virtual environment the effect of a complex tendon-transfer surgery. If muscle synergies only provide a compact description of the regularities in the motor output generated by a non-modular control architecture, compatible and incompatible virtual surgeries should be equally difficult to overcome, because both perturbations can be compensated by appropriately adapting the muscle patterns without any constraint imposed by the synergies. We tested this hypothesis in a virtual environment, in which cursor movements were intuitively controlled in real-time by muscle activity (EMG control). By perturbing the mapping from muscle activity to the force generated at the hand and acting on a virtual mass, visualized as a spherical cursor over a desktop, we could test whether participants were able to adapt their control policies to the new mappings. We showed that, while participants could overcome compatible perturbations, and learn new association between target directions and muscle patterns generated as recombination of existing muscle synergies, they failed to successfully complete the task after incompatible virtual surgeries, despite a reduction of their initial direction errors.

The failure to overcome incompatible virtual surgeries, while predicted by the neural organization of muscle synergies, left several critical questions unanswered. Is it possible to overcome incompatible virtual surgeries by learning new muscle synergies under different conditions, e.g., with enough practice? Which adaptive processes would underlie the acquisition of new synergies? If novel muscle patterns are indeed generated when we learn a new motor skill, it should be in principle possible to learn novel muscle synergies. This could explain the extraordinary muscular independence musicians have (Bengtsson et al. 2005). Transcranial magnetic stimulation (TMS) of the motor cortex of musicians show patterns of joint coordination that cannot be reproduced by analogous stimulation in non-musicians (Gentner et al. 2010). Such a motor independence may be associated with the higher number of muscle synergies available to the musicians. As motor training for as little as 6 weeks can induce changes in white matter (Scholz et al. 2009), skill learning may be related to both the development of new task-specific muscle synergies as well as to changes in the structure or recruitment of existing ones, that is the acquisition of a new synergy-based control policy (Safavynia et al. 2011). However, while it is conceivable that with enough practice novel muscle synergies can be learned (Ericsson et al. 1993), the learning processes underlying the acquisition of novel muscle synergies are not well understood.

During motor adaptation in response to a change in the environment or in the body, a well-known and accurate mapping of goals into motor commands (i.e., a control policy) must be adjusted to reduce performance errors (Krakauer et al. 2019; Shadmehr et al. 2010). This recalibration of motor commands may be achieved either by selecting an alternative goal while using the same control policy or by modifying the control policy. Moreover, adjustments may occur in both feedforward and feedback control mechanisms. While feedback control is necessary for correcting an ongoing movement when an error occurs (Diedrichsen 2007; Diedrichsen et al. 2010; Goodale et al. 1986; Pélisson et al. 1986; Scott et al. 2015), adjusting feedforward control enables to prevent further errors (Shadmehr et al. 2010; Shadmehr and Mussa-Ivaldi 1994; Wolpert et al. 2011). A feedforward control policy based on an inverse model can be learned with a direct inverse modelling approach, where samples of motor commands and movement outputs are used to train the inverse model directly, or with a distal supervised learning approach, where the inverse model is trained indirectly, through the intermediary of a learned forward model of the plant (Jordan 1996). The update of the internal forward model after a perturbation may be driven by the errors in the predicted outcome of a movement (Mazzoni and Krakauer 2006). However, if the desired movement has never been performed before, there is no prior knowledge about the environment of the novel task space. If no internal forward model is available, the motor system might adapt the motor commands needed to successfully perform the desired tasks by training an inverse model as in “goal babbling” (Rohde et al. 2019; Rolf et al. 2010). The main idea of goal babbling is to generate knowledge by exploration by adding some exploratory noise to the motor commands generated according to the existing inverse model and using the error between the motor command predicted by the inverse model as necessary to generate the observed outcome and the actual motor command to train the model. However, in this situation, if the inverse model maps desired outcomes into inappropriate motor commands, it may be impossible to learn the correct motor commands (Sanger 2004).

How learning may occur in a synergy-based control architecture has received much less attention. Adaptation tasks, in which pre-existing muscle synergies can be used to overcome the perturbation, such as after a compatible virtual surgery or a visuomotor rotation, may simply require learning a new mapping between goals and synergy combination coefficients, without having to adjust the synergies. One possibility for learning such a new mapping between goals and synergy combination coefficients is to minimize the motor performance error, i.e., the error between the state of the end effector after the motor command and the desired state. This can be achieved by updating the parameters defining the mapping by gradient descent. As the back propagation of the motor performance error to a motor command error requires a forward model of the plant, it is also necessary to concurrently learn a forward model by minimizing the motor performance prediction error. Thus, combined learning of a forward model and learning of the synergy combination coefficient providing an inverse model would constitute an implementation of a distal supervised learning approach (Jordan 1996; Pierella et al. 2019) in a synergy-based control architecture.

However, such a learning approach cannot succeed in learning an inverse model after an incompatible virtual surgery (Berger et al. 2013). This is because an incompatible virtual surgery makes muscle patterns originally generating a force ineffective, i.e., remaps the muscle patterns onto the null space. Moreover, in the case of incompatible virtual surgeries, no prior information about the novel mapping of muscle activity into the virtual task space is available to the participants. Novel muscle patterns are required not only to perform the correct cursor movements, but any movements outside the one direction determined by the muscle synergies. This pattern has never been generated before and cannot be generated by recombining the subject-specific available synergies. Thus, in a synergy-based control architecture, adaptation to incompatible virtual surgeries (as for learning a new motor skill) requires learning new muscle synergies together with a new control policy that maps goals into muscle patterns. However, it is not clear how such learning processes are implemented.

We hypothesize that, for learning new muscle synergies, it is necessary to explore new directions in muscle space and thereby gain experience about the novel mapping between motor commands and forces affecting the end effector in task space. Therefore, one way to improve performance after incompatible virtual surgeries may be to allow enough time for exploring how new muscle patterns affect cursor movements, i.e., to allow task space exploration. In the present study, we dropped a limitation of our previous experimental protocol, in which the duration of each movement trial was limited to 2 seconds and the time available to relate exploratory motor commands to cursor movements was severely limited. We found that, by allowing 15 seconds to complete each trial, thus providing time for task space exploration, participants significantly improved trial success rate after incompatible virtual surgeries. Improvements in movement direction accuracy after incompatible surgeries occurred faster for corrective movements than for the initial movement, suggesting that learning of new muscle synergies is more effective when used for feedback control. Moreover, reaction time after incompatible virtual surgeries was significantly higher than after compatible virtual surgeries, suggesting an increased use of an explicit adaptive process to overcome incompatible virtual surgeries. These results indicate that, when needed, new muscle synergies can be learned, providing a novel perspective on the exceptional human capacity for acquiring a wide variety of novel motor skills with practice.

## MATERIALS AND METHODS

### General approach

Naive participants were instructed to displace a spherical cursor and reach targets placed on a horizontal plane in a virtual environment displayed stereoscopically on a computer screen. The cursor displacement was controlled according to either the isometric force applied at the hand (force control) or the force estimated from the EMG activity recorded from several shoulder and arm muscles (myoelectric or EMG control). At the beginning of the experimental session, the reaching task was performed under force control. The force and EMG data collected from each participant were then used to estimate an individual mapping of EMG activity patterns onto hand forces (EMG-to-force matrix) by multiple linear regressions, and to identify the muscle synergies by non-negative matrix factorization (NMF). For the rest of the experimental session, participants performed the reaching task using EMG control. During EMG control, each muscle contributes with a force vector in a specific direction to the resultant force vector which causes the displacement of the cursor in a particular direction in the task space (in the virtual environment). Because in EMG control the cursor moves under the effect of the force estimated (in real-time) using the EMG-to-force linear mapping, we can arbitrarily modify such mapping, thus altering the (simulated) force associated to each individual muscle as it would have occurred after a complex tendon-transfer surgery (Berger et al. 2013). For this reason, we refer to the remapping of the forces associated to each muscle as a “virtual surgery”. We could then compare the effects of a compatible virtual surgery with those of an incompatible one. Details of the experimental setup, experimental protocol, and online/offline data analysis (including estimation of the EMG-to-force matrix, identification of muscle synergies, construction of virtual surgeries) are described in (Berger et al. 2013) and are only briefly summarized below.

### Participants

Eight healthy, right-handed human participants (age: 27.75 ± 5.8 years, mean ± SD; 5 females) participated in the experiment after giving written informed consent. All procedures were conducted in conformance with the Declaration of Helsinki and were approved by the Ethical Review Board of Fondazione Santa Lucia (CE/AG4-PROG.222-34).

### Experimental setup

Participants sat in front of a desktop with their right forearm inserted in a splint, immobilizing hand, wrist, and forearm. A 21-inch LCD monitor, displaying a virtual desktop matching the real desktop, occluded the participant’s view of their hand. A steel bar at the base of the splint was attached to a 6-axis force transducer (Delta F/T Sensor; ATI Industrial Automation) recording isometric forces and torques. Surface EMG activity was recorded from 16 muscles acting on the shoulder and elbow joints: brachioradialis (BracRad), biceps brachii long head (BicLong), triceps brachii lateral head (TriLat), triceps brachii long head (TriLong), triceps brachii medial head (TriMed), infraspinatus (InfraSp), anterior deltoid (DeltA), middle deltoid (DeltM), posterior deltoid (DeltP), pectoralis major (PectMajClav), pectoralis major (PectMajStern), teres major (TerMaj), latissimus dorsi (LatDorsi), upper trapezius (TrapUp), middle trapezius (TrapMid), and lower trapezius (TrapLow). EMG activity was recorded with active bipolar electrodes (DE 2.1; Delsys), band pass filtered (20–450 Hz), and amplified (gain 1000, Bagnoli-16; Delsys). Force and EMG data were digitized at 1 kHz using an analog-to-digital PCI board (PCI-6229; National Instruments). Cursor motion was simulated in real time using an adaptive mass-spring-damper (MSD) filter (further details are described in Berger et al., 2013). Moving either under the effect of the actual force recorded by the transducer force (force control) or the force estimated in real time from the recorded and rectified EMGs (EMG control) using a linear mapping (see below, EMG-to-force matrix).

### Experimental protocol

At the beginning of the experimental session, participants performed two blocks of trials in force control. In the first force control block (B1), the mean maximum voluntary force (MVF) along eight directions (separated by 45°) in the horizontal plane was estimated as the mean of the maximum force magnitude recorded across 16 trials in which participants were instructed to generate maximum force in each direction. Participants were then instructed to reach the targets moving the cursor quickly and accurately by applying forces on the splint. At the beginning of each trial (Figure 1*A*), participants were instructed to maintain the cursor within a transparent sphere at the central start position for 1 second (tolerance of 2% MVF). Next, a go signal was given by displaying a transparent target sphere while the start sphere disappeared. Participants were instructed to reach the target as quickly as possible and to remain there for 1 second (tolerance of 2% MVF). After successful target acquisition, the cursor and the target disappeared, indicating the end of the trial. Each trial had to be completed within 15 seconds from its onset. In the second force control block (B2 in Figure 1B), participants performed 72 trials to targets in eight force directions and at three force magnitudes corresponding to 10, 20, and 30% of MVF (randomized order, 3 repetitions for each direction and magnitude). After this block, there was a 5-minute break to process the recorded data and to compute EMG-to-force matrix and synergies necessary to construct a participant-specific myoelectric controller and participant-specific virtual surgeries. After the initial two blocks in force control, for the rest of the experimental session participants performed the task in EMG control. Each EMG control block (Figure 1*B*) consisted of 24 trials with the 8 targets at 20% MVF in random order, with exception of the blocks without visual feedback of the cursor position (HIDDEN), which consisted of 8 trials each. In the first EMG control block (B3), participants familiarized with the new control modality. Then participants performed two series of blocks of compatible and incompatible virtual surgeries, respectively (see below, *Construction of compatible and incompatible virtual surgeries section*). Each series consisted of 15 blocks: 2 baseline blocks, one HIDDEN block, 8 virtual surgery blocks, one HIDDEN block, 2 washout blocks, and one HIDDEN block. In between the two series, participants rested for 10 min; they were also allowed to rest at any time during the experiment (Figure 1B).

**Figure 1.**
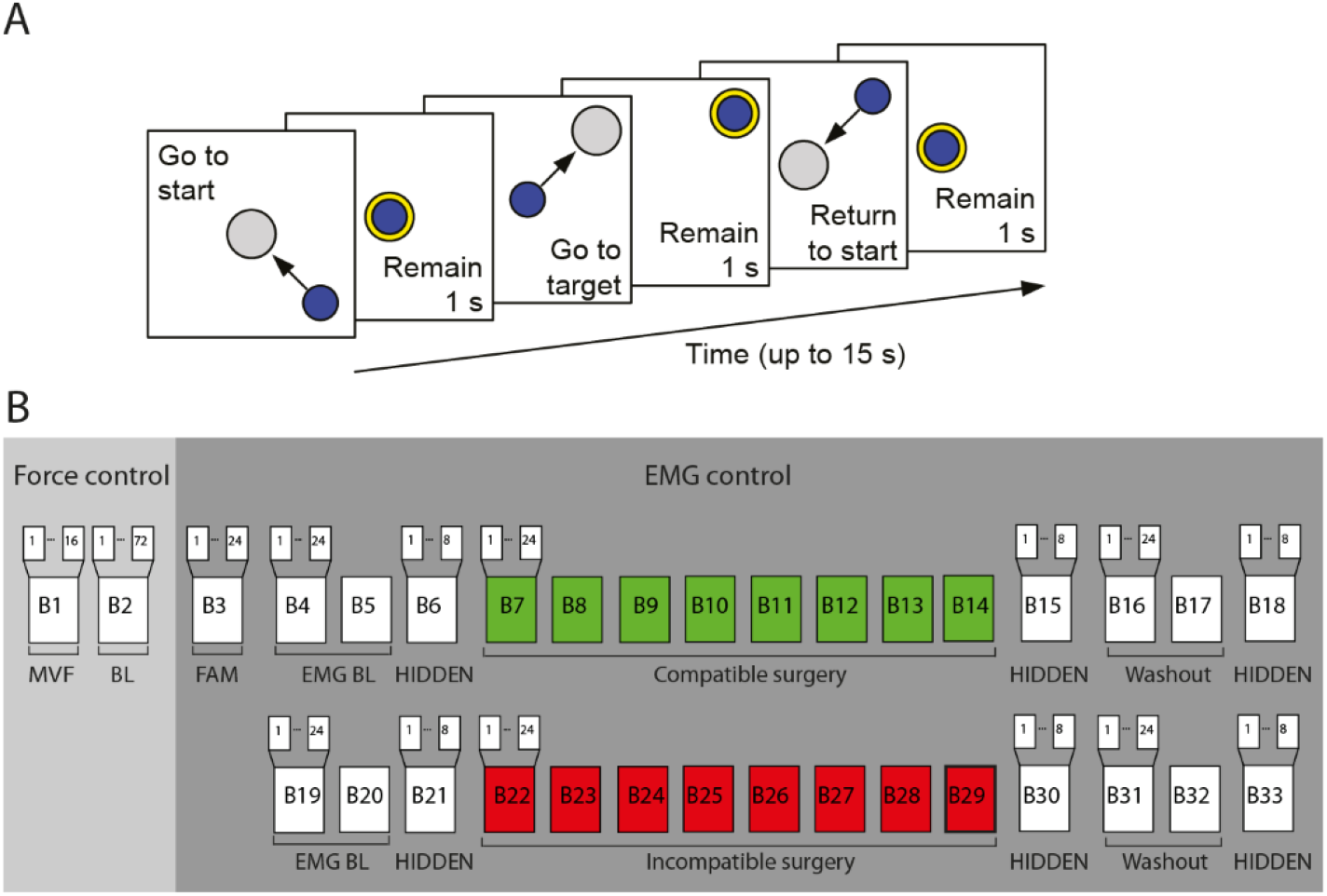
Experimental task and protocol. (A) Participants were instructed to perform a center-out reaching task with a cursor in a virtual environment by applying isometric force on a simulated mass-spring-damper system. They had to maintain the cursor in a central start location for 1 s, reach a target as soon as it appeared at one of 8 peripheral locations, and maintain the cursor at the target for 1 s. (B) Each participant performed a single experimental session consisting of 16 trials of maximum voluntary force generation in 8 directions (MVF), 72 trials of reaching to targets in 8 directions at three force magnitudes (10%, 20%, 30% MVF) in force control, and the following blocks of 24 trials each in EMG control: one block of familiarization, two series of 15 blocks for each surgery type (2 baseline blocks, 1 HIDDEN block, 8 surgery blocks, 1 HIDDEN block, 2 washout blocks, 1 HIDDEN block).

**Table 1.**

| *Subject ID* | *No. of synergies* | *Synergy reconstruction $R^2$* | *I diff (Tc)* | *I diff (Ti)* |
| --- | --- | --- | --- | --- |
| 1 | 6 | 0.92 | 1.36 | 1.35 |
| 2 | 4 | 0.92 | 1.33 | 1.32 |
| 3 | 5 | 0.90 | 1.20 | 1.20 |
| 4 | 5 | 0.92 | 1.61 | 1.59 |
| 5 | 6 | 0.92 | 1.27 | 1.26 |
| 6 | 7 | 0.92 | 1.60 | 1.58 |
| 7 | 4 | 0.93 | 1.32 | 1.32 |
| 8 | 4 | 0.92 | 1.05 | 1.05 |

### EMG-to-force mapping

For EMG control, we created a mapping between recorded muscle activity and the movement of a computer cursor. The force generated at the hand was approximated as a linear function of the activation of muscles acting on shoulder and elbow: **f** = **H m**, where **f** is the generated two-dimensional force vector, **m** is the 16-dimensional vector of muscle activations, and **H** is a matrix relating muscle activation to force (dimensions 2 × 16). Such EMG-to-force matrix (**H**) was estimated using multiple linear regressions of each applied force component, low-pass filtered (second-order Butterworth; 1 Hz cut off), with the EMG signals recorded during the initial force control block (static phase, i.e., time from target acquisition until target leave), low-pass filtered (second-order Butterworth; 5 Hz cut off) and normalized to the maximum EMG activity during the generation of MVF. Only muscles which significantly contributed to the EMG-to-force mapping, were included for the calculation of the EMG-to-force mapping. This was estimated by the confidence interval (95%) for the coefficient estimates of the multiple linear regression. Figure 2*A* illustrates the force associated with each muscle (the columns **h_i_** of the EMG matrix) estimated in participant 1.

**Figure 2.**
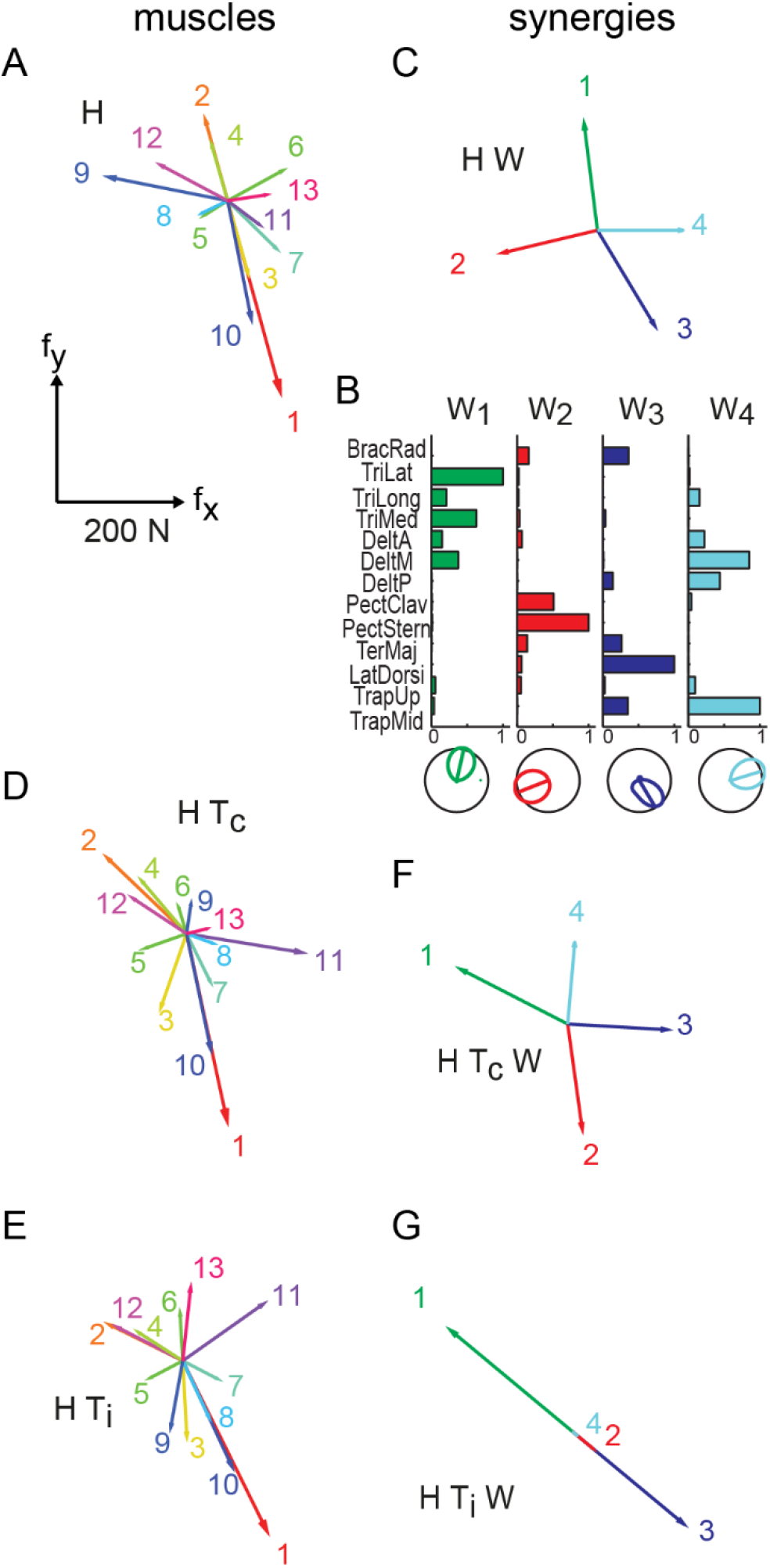
Examples of EMG-to-force matrix, synergies, and virtual surgeries. (A) EMG-to-force matrix **H** estimated for participant 1 from EMG and force data recorded during the generation of planar isometric forces. Each column of **H**, representing the planar force generated by one muscle, is illustrated by a coloured arrow (1: brachioradialis, 2: triceps brachii lateral head, 3: triceps brachii long head, 4: triceps medial head, 5: anterior deltoid, 6: middle deltoid, 7: posterior deltoid, 8: pectoralis major clavicular, 9: pectoralis major sternal, 10: teres major, 11: latissimus dorsi, 12: upper trapezius; 13: middle trapezius). (B) Muscle synergies (matrix **W**) are identified by NMF from the EMG data. Each column of **W**, a vector specifying a specific pattern of relative level of muscle activation, is illustrated by color-coded horizontal bars. Below the respective polar plot of the directional tuning of the four synergies. (C) Forces associated to each muscle synergy (i.e., columns of the matrix product **H W**) span the entire force space. (D) Forces generated by muscles after a compatible virtual surgery obtained by recombination of the original forces as after a complex re-arrangement of the tendons (**T_c_**). (E) Muscle forces after an incompatible surgery generated by a rotation matrix (**T_i_**) that maps a vector in the column space of **W** into a vector in the null space of **H**. (F) Synergy forces after the compatible surgery still span the force space. (G) Incompatible rotations align forces associated to all synergies in the same direction, thus synergy forces after an incompatible surgery do not span the entire force space.

### Muscle synergy extraction and selection of the number of muscle synergies

Muscle synergies were identified by NMF (Lee and Seung, 1999) from EMG patterns recorded during force control in the time interval in which the cursor was within the target (static phase): **m** = **W c**, with **W** the *M* × *N* synergy matrix, **c** the *N*-dimensional synergy activation vector, where *N* is the number of synergies, and *M* the number of muscles. EMG patterns were first low-pass filtered (second-order Butterworth filter; 5 Hz cut-off frequency) and rectified, their baseline noise level was then subtracted, and finally normalized to the maximum EMG activity of each muscle recorded during the generation of MVF. The number of synergies *(N)* was then selected according to two criteria: (1) the smallest *N* explaining at least 90% of the data variation; (2) the *N* at which there was a change in the slope of the curve of the *R*^2^ value as a function of *N*. In case of mismatch between the two criteria, we used a third criterion that selected the set of synergies with preferred directions (the direction of the maximum of the cosine function best fitting the directional tuning) of the synergy activation coefficients distributed more uniformly and with the smallest number of similar preferred directions. We thus arranged the preferred direction vectors on a unit circle, considering all adjacent pairs, and we selected the set of synergies with the smallest number of pairs with an angular difference below 20°. Figure 2*B* illustrates the synergies extracted from participant 1, and Figure 2*C* the forces associated with those synergies by the EMG-to-force matrix (**f_i_** = **H w_i_**) estimated in the same participant (Figure 2*A*).

### Construction of compatible and incompatible virtual surgeries

EMG control allows to modify the EMG-to-force map and thereby to perform simulated virtual surgeries on the musculoskeletal system. By virtually rearranging the tendons, we constructed altered mappings that were either compatible or incompatible with the participants’ muscle synergies. For incompatible virtual surgeries, the mapping was altered in a such way that it could not be compensated by recombining the synergies. Compatible virtual surgeries, in contrast, could be compensated by recombining the synergies. We constructed both types of virtual surgeries altering the EMG-to-force mapping (**H**) with a rotation in muscle space: **H′** = **H T,** where **H’** is the transformed EMG-to-force mapping after the virtual surgery and **T** is a rotation matrix (Figure 2*D*, *E*). For each participant, compatible and incompatible virtual surgeries were constructed according to the muscle synergies identified in initial force control block at the beginning of the experimental session. Compatible rotations (**T_c_**) were chosen such that the forces associated to the muscle synergies after the rotation still spanned the whole force space. In contrast, incompatible rotations (**T_i_**) were chosen such that the forces associated to the muscle synergies did not span the whole force space after the rotation. This was achieved by rotating a vector (**w**) of the muscle synergy subspace not in the null space of **H** onto a vector **n** in the null space of **H** not in the synergy subspace. Thus, while both types of virtual surgeries allowed the generation of any planar force with a new muscle activation vector, **m′**, only for compatible virtual surgeries all force directions could be generated by recombining the existing muscle synergies (Eq. 3, **c′**; Figure 2*F*). Incompatible virtual surgeries, in contrast, were constructed such that muscle activation vectors obtained by muscle synergy combinations could only generate forces in one dimension (Figure 2*G*). Further details on the construction of compatible and incompatible virtual surgeries can be found in (Berger et al. 2013).

The compatible rotation angle was chosen such that a difficulty index (I_diff_), defined as the average change across muscles and force targets in muscle activity required to perform the task after the virtual surgery, was similar in the two cases. Specifically, the difficulty index was defined as follows: 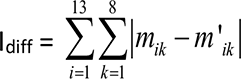, where m_ik_ is the activity of the i^th^ muscle for the k^th^ force target, normalized to the maximum across force targets before the virtual surgery, and m′_ik_ the same activity after the virtual surgery. For each force target k, the muscle activation vectors m_k_ and m′_k_ were computed as the minimum norm, non-negative solution of the equations f_k_ = H m_k_ and f_k_ = H T m′_k_, respectively.

### Data analysis

We quantified the task performance by the percentage of unsuccessful trials, the initial direction error, the direction error after the first movement correction, exploration time, the number of speed peaks, and the length and area of the convex hull of the cursor path. A *successful trial* was defined as a trial in which the cursor reached and remained within the target for the instructed time interval. The *initial direction error* (iDE) was defined as the angular deviation of the initial movement direction of the cursor with respect to the target direction. The iDE was computed as the absolute angle between the direction from the cursor position at movement onset and the target and the direction from the cursor position at movement onset and the cursor position at the first velocity peak. For determining the movement onset, we used a threshold based on the cursor velocity during the initial hold phase, i.e. the interval from 0.5 s before the target go signal to the target go signal, for each trial. The movement onset was defined as the time at which the cursor velocity crossed the trial specific velocity threshold, which was set to five time the maximum velocity during the initial hold phase. The *direction error after the first movement correction* (cDE) was defined as the angular deviation of the movement direction at the time of the second velocity peak with respect to the target direction at that time. The cDE was computed as the absolute angle between the direction from the cursor position at the time of the first trough of the cursor velocity that occurred after the first velocity peak and the cursor position at the time of the second velocity peak and the direction of the cursor displacement at the first subsequent peak velocity. *Exploration time* was defined as the time elapsed from movement onset until the target was reached. If the target was not reached, exploration time was defined as the time from movement onset until the end of the trial. To estimate the *number of speed peaks*, the cursor trajectories were first filtered (second-order Butterworth; 3 Hz cut off) and then the number of velocity peaks were evaluated on the cursor’s velocity during the exploration time (using Matlab function *findpeaks*). The *length of the cursor path* and the *area of the convex hull* of the cursor path (using Matlab function *convexHull*) were evaluated during the exploration time as the smallest convex set that contains the cursor path.

We quantified the changes in the synergistic organization of the muscle patterns throughout the experiment by reconstructing the muscle patterns of each block as combination of the synergies extracted at the initial force control block and computing a reconstruction *R*^2^ value for each block. The reconstruction of the muscle patterns of each block was performed using the NMF algorithm initialized with the synergies extracted from the initial force control block and updating only to the combination coefficients (d’Avella et al. 2006).

We estimated the *reaction time* (RT) based on two different measures. The first measure (RT-cursor) was based on the time of cursor movement onset after the go signal. The second measure (RT-EMG) was based on the onset of muscle activity after the go signal. The threshold based on the onset of muscle activity was defined as the first time after the movement go signal at which any of the recorded muscle activities crossed a muscle and trial specific threshold. To determine the threshold for muscle and trial, we estimated the mean plus two times the standard deviation of the activity of each muscle (low-pass filtered -second-order Butterworth; 5 Hz cut off-and normalized to the maximum EMG activity during the generation of MVF) in the 0.5 s interval prior to the go signal. For each trial, we only considered those muscles whose maximum activity in the entire trial was higher than the mean plus two times the standard deviation of the activity during the movement period in the baseline condition, i.e., the averaged muscle activity in blocks 4 and 5. Movement period was defined as the interval from the time at which the cursor exited the initial sphere until the time the cursor entered the target, or end of the trial in case of an unsuccessful trial.

### Statistical analysis

For the statistical analyses we used linear mixed-effects (LME) models, with exception of the trial success, a binomial measure, for which we used a generalized linear mixed-effect (GLME) model implemented in Matlab 2020b Statistics Toolbox. All analyses included *block id* and *surgery type* as fixed effects, and participant as a random effect, resulting in the following model: *Dependent variable* ∼ *ID* + *condition* + (1 | s*ubject*)). Differences in performance measures between individual blocks were assessed with a t-test after verifying the normally of the data distribution (according to a Lilliefors test).

## RESULTS

Eight healthy participants were instructed to perform a center-out reaching task in a virtual environment by applying isometric forces to a force transducer attached to a hand-wrist splint (Figure 1; see *MATERIALS AND METHODS* for details). After the initial force control blocks, in which forces applied determined the cursor motion in the horizontal plane, the cursor was controlled by EMG signals recorded from up to 16 arm and shoulder muscles (EMG control) throughout the rest of the experiment. The initial force control block allowed to estimate a linear mapping between muscle activity and force at the hand (muscle-to-force matrix) then used to control the cursor intuitively in the EMG control blocks. Thus, during baseline EMG control, each muscle generated a force at the hand in a specific direction in the task space in the virtual environment (Figure 2A). Consistently with our previous results (Berger et al. 2013; Berger and d’Avella 2014) all participants performed well during the baseline EMG-control session, i.e., they managed to move the cursor along approximately straight paths from a central start location to one of eight targets arranged on a circle (Figure 3, *first column*), with small initial direction error of 14.5 ± 7.9 degrees (mean ± SD across participants, familiarization block). Success rates during this baseline blocks were high with an average trial success rate of 98.4 ± 3.1% (mean ± SD, familiarization block).

**Figure 3.**
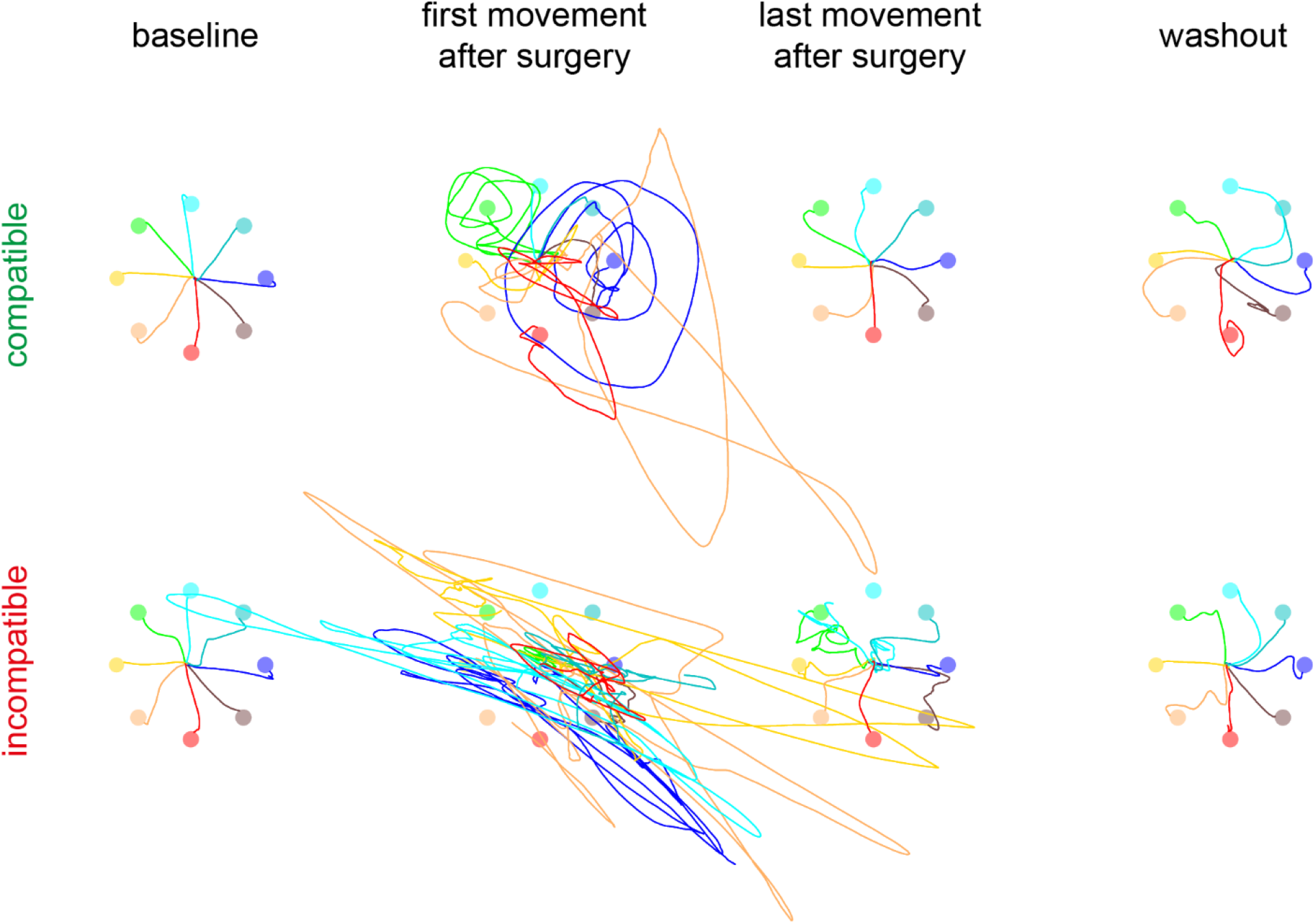
Example of cursor trajectories. Trajectories of the cursor on the horizontal plane during individual trials of participant 1 are shown for different targets (*colour coded*), before undergoing a virtual surgery (baseline, *first column*), immediately after a virtual surgery (*second column*), at the end of the exposure to the virtual surgery (*third column*), and after undoing the virtual surgery (washout, *last column*), for compatible (*first row*) and incompatible (*second row*) virtual surgeries. The cursor motion was simulated in real-time as that of a mass attached by a damped spring to the center of the real handle under the force applied to the handle estimated from the recorded EMGs (see *Materials and Methods*). Trajectories are shown from target *go* signal until the target was reached and in case of unsuccessful until the end of the trial.

After the baseline EMG-control session the mapping between muscle activity and cursor movement was altered by performing virtual surgeries. The direction of the force generated by each muscle changed (Figure 2D and E) such that the forces generated by the muscle synergies still spanned the whole plane after a compatible surgery (Figure 2F) but were aligned in a single direction after an incompatible surgery (Figure 2G). Immediately after a virtual surgery, irrespective of its type, cursor movements were poorly controlled by the participants. Indeed, the muscle patterns normally used to generate a force towards the target now directed the cursor in a different direction (Figure 3, *second column)*. However, in case of a *compatible* virtual surgery (Figure 2, *second column* top), cursor trajectories showed a curved path with progressively smaller radii around the target position as trial time increases. After the *incompatible* virtual surgeries, in contrast, the cursor trajectories were aligned along a single direction, which matched the synergy force direction (compare the synergy-to-force mapping shown for the same participant in Figure 2G for the incompatible virtual surgery). As expected, at the end of the compatible perturbation blocks, the participant, as illustrated in Figure 2, could easily reach all targets. Surprisingly, however, after the incompatible perturbations, while initially the performance was poor, with low trial success, at the end of the incompatible blocks the same participant managed to reach all but one target (Figure 3, *third column)*. After training each virtual surgery for 8 blocks of 24 trials, i.e., for a total of 192 trials, the virtual surgery was removed and the muscle-to-force mapping was restored to the original, intuitive, baseline mapping. Immediately after removing the compatible virtual surgery, an after-effect was present as the cursor paths showed large curvatures deviating from a straight line in the opposite direction than in the first perturbed block (Figure 3, *fourth column top)*. An after-effect was less prominent after removal of the incompatible virtual surgery (*bottom*), where most trajectories showed straighter paths to the targets.

### Participants adapted to compatible virtual surgeries faster than to incompatible virtual surgeries

To assess task performance, we estimated the angular error of the cursor initial direction (Figure 4A). The iDE was significantly larger in the first block after the virtual surgery than in the last block preceding it, for both compatible and incompatible virtual surgeries (p < 10^-4^, t-test). With practice, participants improved task performance, as they improved their ability to generate appropriate muscle patterns (significant block effect for both surgeries, p < 10^-4^ and p = 0.0015 for compatible and incompatible virtual surgeries, respectively). However, the iDE was significantly larger at the end of incompatible virtual surgery than at the end of the compatible virtual surgery (p < 10^-4^, t-test, comparing the last block of each virtual surgery). This result revealed a faster adaptation to the compatible than to the incompatible virtual surgery, confirming our previous results (Berger et al. 2013).

**Figure 4.**
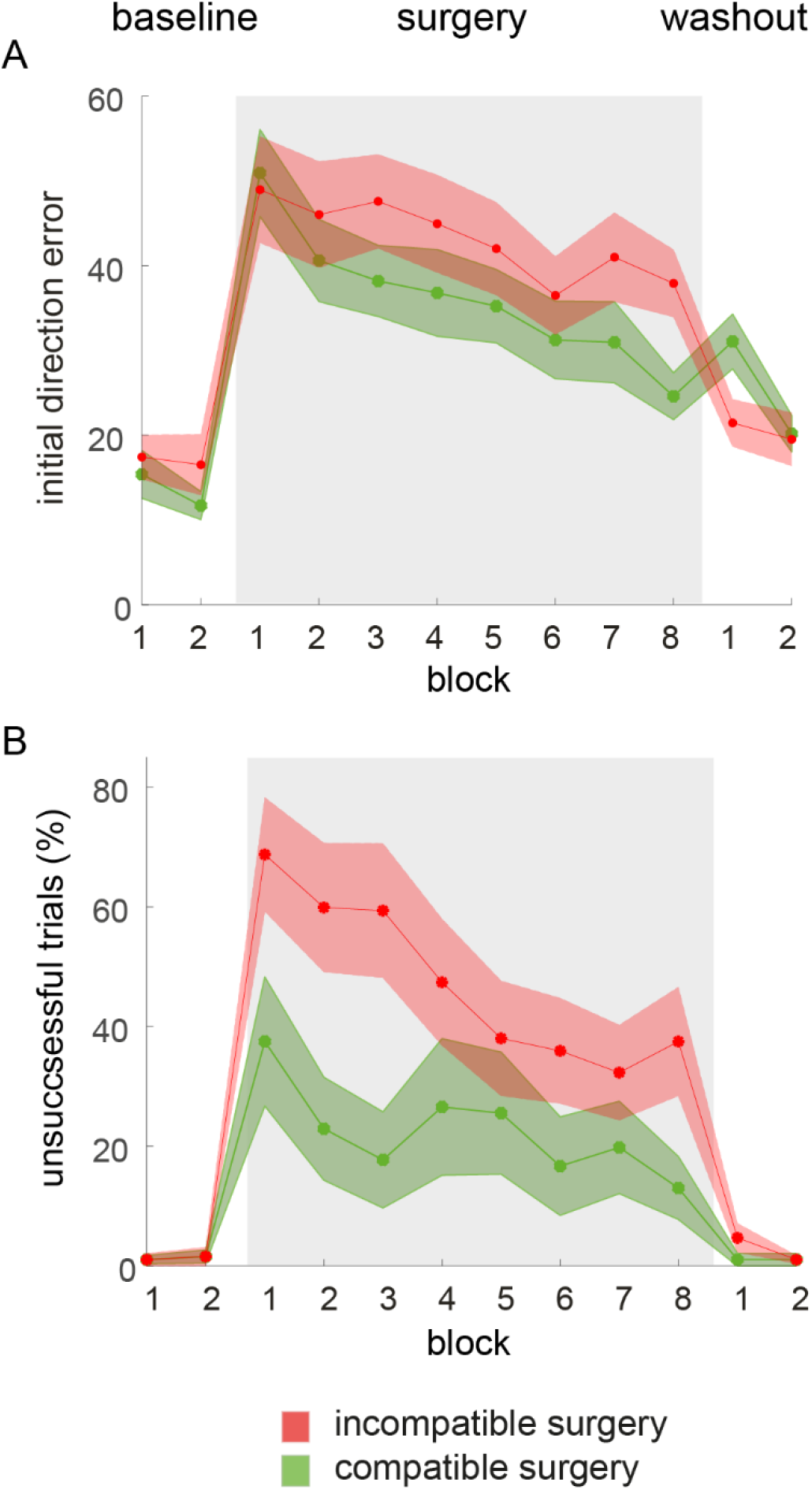
Comparison of task performance changes, assess by initial direction error and unsuccessful trial rate, after virtual surgery. (A) Direction error of the initial movement direction with respect to the target direction, averaged across participants and blocks of 24 trials (solid lines, shaded areas indicate SE) for compatible virtual surgery (*green*), incompatible virtual surgery (*red*). Differences between compatible and incompatible conditions are significant at the end of the exposure to the perturbation (surgery block 8) in the virtual surgery experiments. (B) Fraction of trials in which the cursor did not reach and hold on the target, averaged across participants and blocks of 24 trials (solid lines, shaded areas indicate SE) for compatible virtual surgery (*green*), incompatible virtual surgery (*red*). Differences between compatible and incompatible conditions are significant at the end of the exposure to the perturbation (surgery block 8) in the virtual surgery experiments.

### After-effects were larger after compatible virtual surgeries than after incompatible virtual surgeries

To further assess the performance differences after compatible and incompatible virtual surgeries we considered the after-effects in the first washout blocks, when the EMG-to-cursor mapping was restored to the original, intuitive, baseline mapping. Compatible and incompatible surgeries showed different after-effects (Figure 4A), with a significantly higher iDE after the compatible surgery than after the incompatible surgery (p < 10^-4^, t-test). The iDE in the first washout block after compatible virtual surgery was significantly higher than the error in the last baseline block before the surgery (p < 10^-4^, t-test) indicating that learning had occurred (Shadmehr et al. 2010). On the contrary, the iDE of the first washout block after the incompatible virtual surgery was not significantly higher than the error in the baseline block before the surgery (p = 0.2344, t-test) and even significantly lower than the error in the last incompatible surgery block (p = 0.0221, t-test), indicating an immediate return to baseline behaviour.

### Participants increased trial success rate after incompatible virtual surgeries

To assess the effects of the entire trial duration, which was longer than in our previous protocol (Berger et al. 2013), we computed trial success rate (Figure 4B). Differently from the iDE, which provides a measure of acquisition of a novel control policy involved in the feedforward generation of the initial muscle pattern, trial success also considers contribution of online error corrections throughout the trial due to visual feedback. A trial was classified as successful if the participant could both reach the target within 15 seconds and remain therein for 1 second. For both surgery types, the fraction of unsuccessful trials was significantly larger in the first block after the surgery than in the last block preceding it (p = 0.0125 after compatible and p < 10^-4^, t-test, after incompatible surgery). With practice, participants decreased the fraction of unsuccessful trials not only after the compatible virtual surgery (significant block effect, p < 10^-4^, GLME) but, interestingly and most importantly, also after the incompatible surgery (significant block effect, p < 10^-4^, GLME). Thus, participants could learn to control the cursor and to guide it successfully to the target even after the incompatible virtual surgery. Nevertheless, performance was significantly worse at the end of incompatible surgery than at the end of the compatible surgery (p = 0.0074, t-test).

### Compatibility of muscle patterns after surgeries to the muscle synergies

Since performance improved after incompatible surgeries, even if performance was significantly worse at the end compared to that of compatible surgeries for both trial success and direction error, we assessed whether this result was associated with changes in the muscle patterns that could not be captured by the original muscle synergies. This would indicate that performance improvements were due to exploration of muscle activation space and generation novel coordination patterns. To this purpose, we quantified whether the muscle synergies identified in the initial force control block, i.e., before any virtual surgery was introduced, could reconstruct the muscle patterns observed after the virtual surgeries. During the baseline blocks and immediately after a virtual surgery, we expected participants to activate their existing muscle synergies to move the cursor. Therefore, a high similarity between actual muscle patterns and their reconstruction using muscle synergies should be revealed. For the same reason, we also expected a high similarity at the end of the learning period of the compatible virtual surgery. Indeed, while compatible surgeries could be overcome simply using the existing muscle synergies and re-associating them with new movement directions, incompatible virtual surgery could be compensated only by generating novel muscle patterns. In the latter case we therefore expected participants to either use new muscle activity patterns or not be able to perform any cursor movements and remain in the one-dimensional task space determined by the perturbed muscle synergy forces.

Figure 5 shows a representative example of muscle patterns for different trials to one target (direction 90 degrees) throughout an experimental session, as well as the corresponding reconstructions obtained with the muscle synergies (red lines). The muscle synergies accurately reconstructed the muscle patterns during the intuitive, baseline blocks (Figure 5, B5, B17 with *R*^2^ of 0.81 and 0.75, respectively), as well as at the beginning and at the end of the exposure to the compatible virtual surgery (B6 and B13, with *R*^2^ of 0.8 and 0.7, respectively), and immediately after the introduction of the incompatible virtual surgery (B18, *R*^2^ = 0.68), with *R*^2^ values ranging from 0.73 to 0.81. However, at the end of the exposure to the incompatible virtual surgery the muscle patterns could not be reconstructed any longer by the initial muscle synergies (*R*^2^ = 0.3). A closer inspection of the corresponding cursor paths shows that the participant was not able to reach the target right after the incompatible surgery (light magenta), whereas at the end of the incompatible surgery period the participant was able to reach the target (dark magenta). On average across all participants, we found a significant reduction of the muscle pattern reconstruction quality at the end of the exposure to incompatible surgeries (Figure 6, significant block effect, p < 10^-4^, LME) but not after compatible virtual surgeries (p = 0.603, block effect, LME).

**Figure 5.**
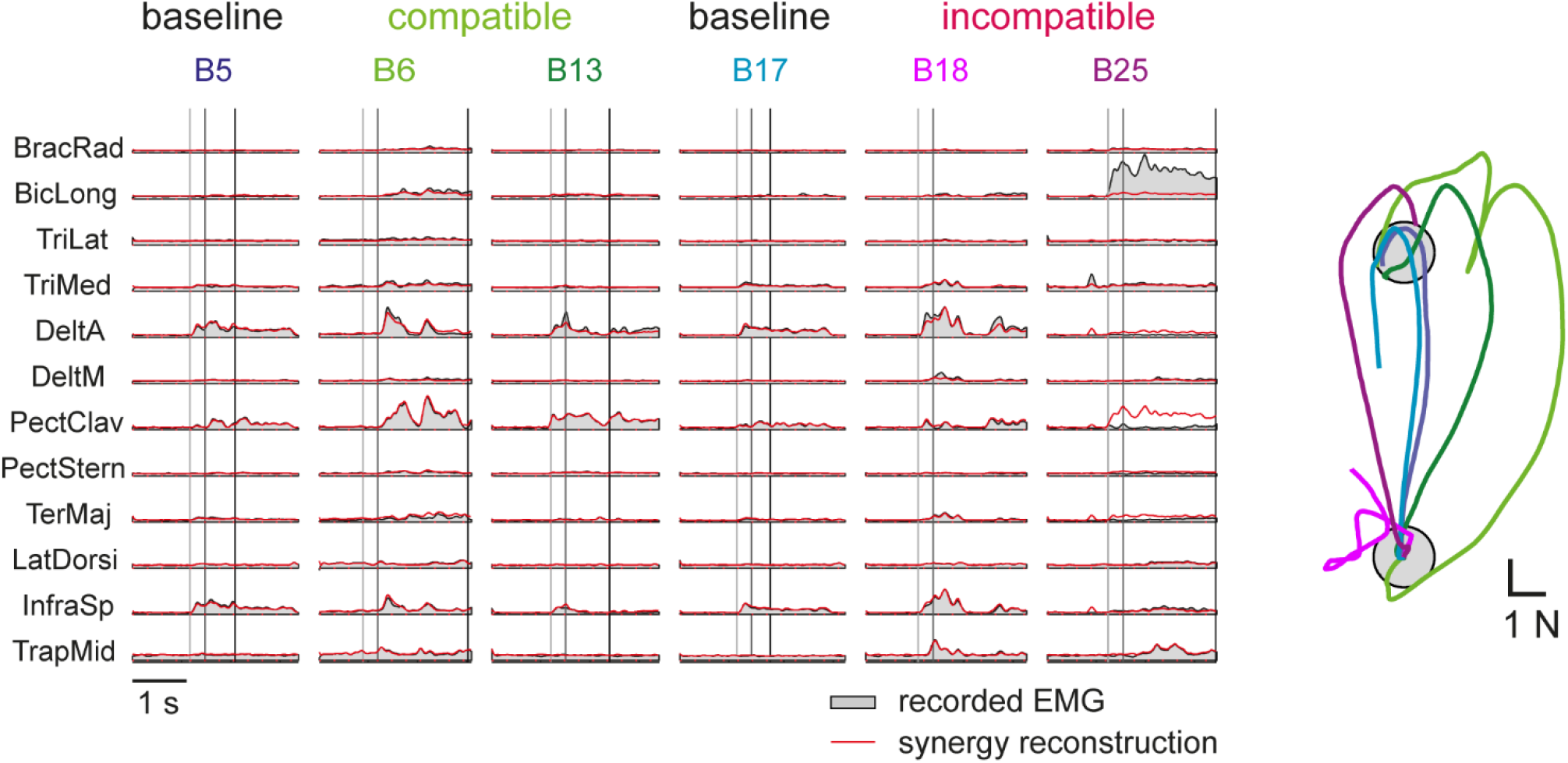
Examples of muscle pattern reconstruction by baseline synergies. (A) Muscle patterns recorded in participant 4 for different trials (*different columns*) to one target (direction 90 degrees, *grey areas*) throughout an experimental session and their reconstruction by the synergies extracted from the initial baseline block (*red lines*). The vertical lines indicate the time of movement onset (1), the time of the first peak of the cursor tangential velocity (2), and the time of target acquisition (3). The corresponding cursor trajectories and events (1-3) are shown for each trial above the muscle activation waveforms.

**Figure 6.**
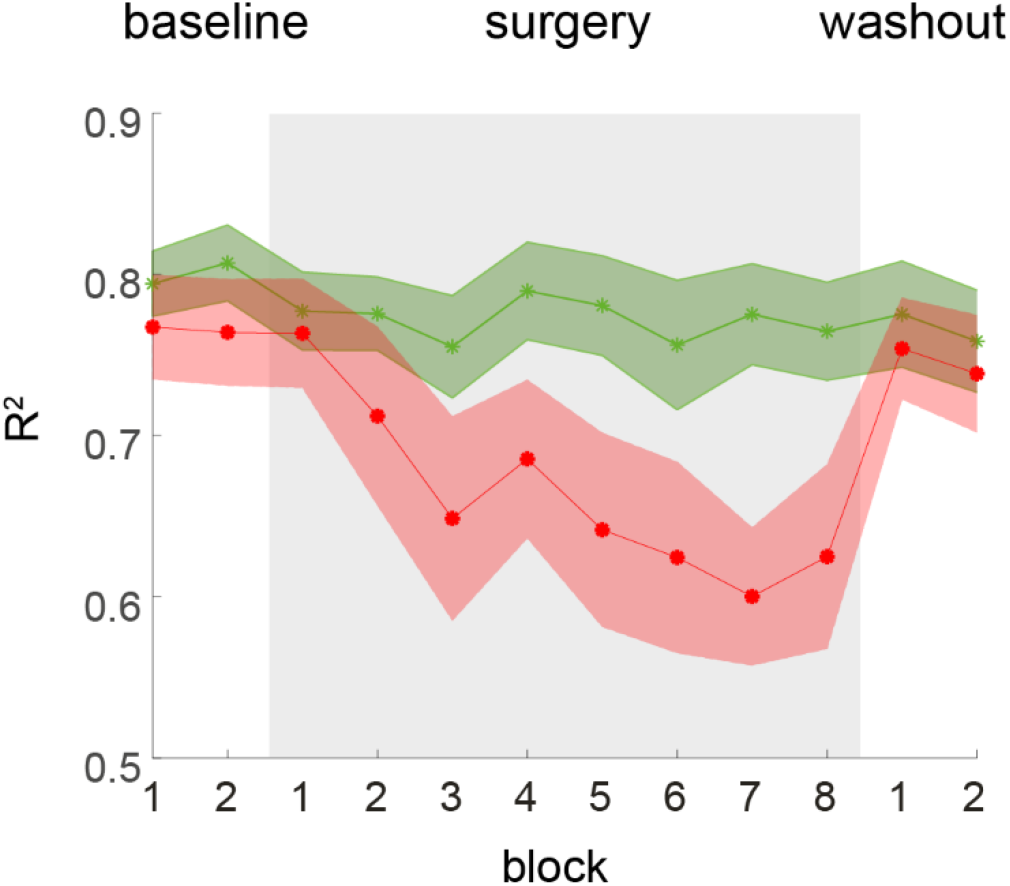
Reconstruction quality of muscle patterns by synergy combinations after virtual surgeries. Synergy reconstruction error (R^2^) averaged across participants and blocks of 24 trials (solid lines, shaded areas indicate s.e.m.) for compatible surgery (*green*) and incompatible surgery (*red*). The reconstruction quality is significantly reduced only during the exposure to incompatible surgeries, indicating a re-organization of the muscle patterns.

In sum, to overcome a compatible virtual surgery, participants generated muscle patterns by recombining the existing muscle synergies normally used to perform the task and re-associated them with new cursor movement directions. In contrast, to overcome an incompatible surgery, since the set of muscle synergies usually employed to perform the task became ineffective, participants generated novel muscle patterns that could not be obtained recombining existing synergies.

### Participants explored the novel task space after virtual surgeries using visual feedback

Additional task performance measures, such as number of movement corrections and the convex hull of the trajectory may shed light into how visual feedback is used to explore the task space and to guide cursor movements towards successful target acquisition. As the maximum trial duration was 15 seconds, participants could explore the effects of novel muscle activation patterns on cursor movements and use visual feedback to correct cursor movement. The total time participants spent trying to reach the target across all blocks was 16.6 ± 6.4 minutes for the compatible virtual surgery and 25.1 ± 8.6 minutes for the incompatible virtual surgery. This is the time in which participants explored the task space. For both virtual surgeries, the target acquisition time per trial, i.e., trial time, decreased significantly over the duration of the experiment (Figure 7A, significant block effect for both surgeries p < 10^-4^, LME). However, participants were faster already in the first block after compatible surgeries than in the first block after incompatible surgeries (p = 0.0036, t-test). This performance reflects a better control of the cursor immediately after surgery onset and is in accordance with the higher trial success rate we observed immediately after compatible virtual surgery. The area spanned by the cursor trajectories, quantified by the convex hull of the path of the cursor in each trial, was smaller after incompatible than after compatible virtual surgeries (Figure 7B), although the path length was higher for the former than for the latter surgeries (Figure 7C). The fact that immediately after the incompatible virtual surgery a longer path length was associated to a smaller convex hull indicates that participants initially (i.e., immediately after the incompatible surgeries) did not explore the whole task space and remained mostly confined along the synergy force direction.

**Figure 7.**
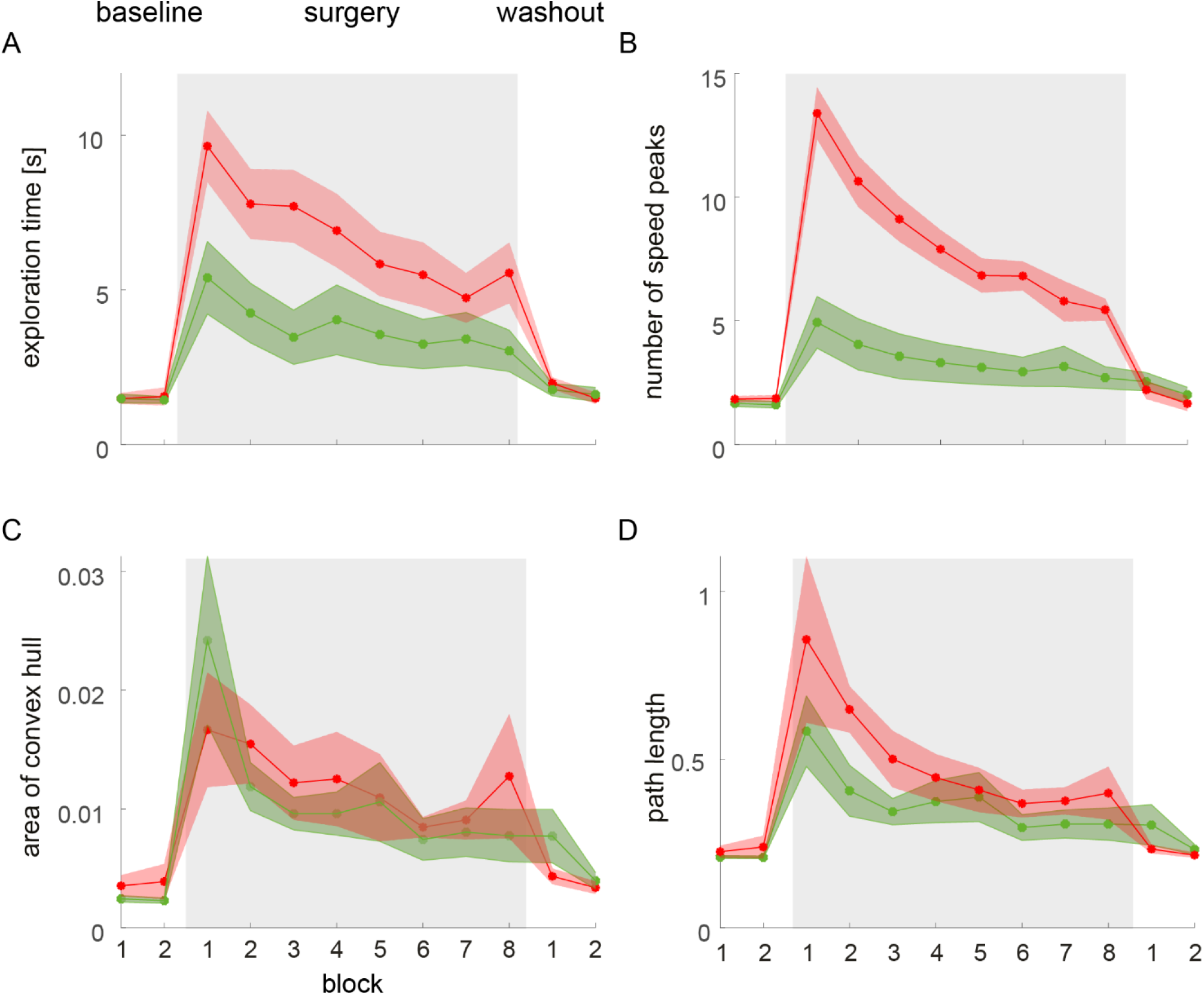
Exploration after virtual surgeries. (A) Time participants need to reach the target, averaged across participants and blocks of 24 trials (solid lines, shaded areas indicate s.e.m.). (B) Numbers of speed peaks, averaged across participants and blocks of 24 trials. (C) Area of convex hull of the cursor path, averaged across participants and blocks of 24 trials. (D) Cursor path length, averaged across participants and blocks of 24 trials.

A closer inspection of the characteristics of cursor trajectories revealed that the number of peaks in the speed profile increased significantly after both surgeries, indicating that participants made use of visual feedback to correct the cursor movements (Figure 7D). The number of velocity peaks increased significantly in the first perturbed block with respect to the last baseline block for both perturbation types (p < 10^-4^, t-test, compatible and incompatible surgeries, respectively), and then decreased significantly during the perturbation blocks after both surgeries with perturbed block number (significant block effect for both surgeries, p < 10^-4^, LME). The presence of multiple online corrections, together with the significant improvement of trial success, indicates that participants were able to use visual feedback to improve their control of the cursor after the incompatible perturbation. At the end of each surgery types the number of peak speeds was significantly higher than during baseline, indicating that also after the compatible virtual surgery online corrections had not returned to baseline levels, however with an average value of 2.7 compared to 5.4 corrections after the incompatible virtual surgery, the number of corrections was significantly lower (p = 0.0091, t-test).

In sum, participants decreased their target acquisition time as well as the number of corrective movements after both surgery types, suggesting that they improved feedback control throughout the virtual surgery period to guide and correct cursor movements. These results indicate that visual feedback together with task space exploration is important for successful task performance for both virtual surgery types.

### Effect of feedback mechanisms on the direction error after virtual surgeries

We have shown above that the iDE significantly improved after incompatible virtual surgeries (section *Participants adapt to compatible virtual surgeries faster than to incompatible virtual surgeries*). The iDE depends on muscle patterns that are activated before visual feedback is available, thus reflects inaccuracy in feedforward control. However, since participants made use of online corrections during task space exploration, we wondered which effect these had on the direction error of corrective movements. We therefore analyzed the direction error of the first movement correction (cDE) during the exposure to a virtual surgery. Figure 8 shows the iDE and cDE, for compatible (Figure 8A) and incompatible virtual surgeries (Figure 8B). In the first block after a compatible virtual surgery, the cDE was significantly lower than the iDE (p = 0.0077, t-test) and then remained at the same level throughout the whole surgery period (with no significant effect of block p = 0.6517, LME). Thus, participants were immediately able to generate accurate online corrections that led to lower direction errors than in the initial movement. With practice participants improved significantly only their iDE (significant effect of block, p < 10^-4^, LME), however with no further improvement of cDE. These results show that participants were immediately able to use effectively feedback control however with practice they significantly improved the accuracy of feedforward control.

**Figure 8.**
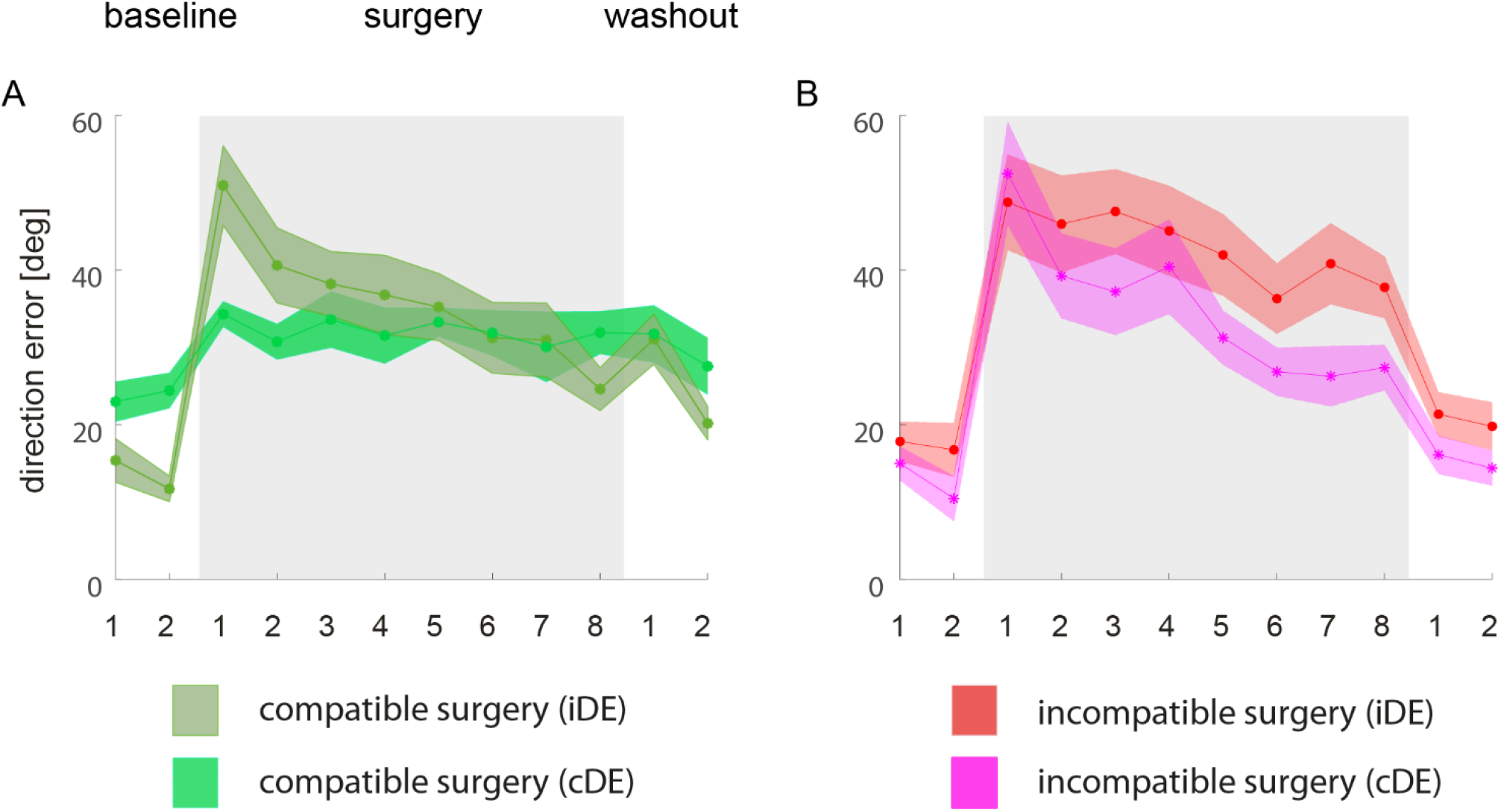
Comparison of direction error after virtual surgeries for the initial and the corrected movement. (A) Angular error of the initial movement direction (green) and first corrected movement (light green) averaged across participants and blocks of 24 trials (solid lines, shaded areas indicate s.e.m.) for compatible virtual surgeries (*green*). (B) Angular error of the initial movement direction (red) and first corrected movement (magenta) averaged across participants and blocks of 24 trials (solid lines, shaded areas indicate s.e.m.) for incompatible virtual surgeries.

In contrast, in the first block after an incompatible virtual surgery, the cDE was comparable to the iDE (p = 0.38, t-test), but then decreased significantly faster than the iDE (p < 10^-4^, LME) and was significantly lower than the iDE at the end of the incompatible perturbation (p = 0.010, t-test). Thus, during the incompatible perturbation participants not only significantly improved the directional error of the first corrective movement (significant effect of blocks, p < 10^-4^, LME) but they also improved it significantly faster than the directional error of the initial movement. These results suggest that participants, after an incompatible surgery, although they improved the accuracy of feedforward control, were more effective to improve feedback control.

### Increased reaction time after incompatible virtual surgeries

Reaction time (RT) increased with respect to baseline performance according to two different measures (Figure 9), both after compatible virtual surgeries (p = 0.0022 and p = 0.0003, LME, for the onset of cursor movements and the onset of muscle activity, respectively) and after incompatible virtual surgeries (p < 10^-4^, LME, for both measures). RT was significantly longer after incompatible virtual surgeries than after compatible virtual surgeries (p < 10^-4^, LME, for both measures of RT). At the end of the incompatible virtual surgery RT was still significantly higher than during baseline for both measures of RT (p = 0.046 and 0.040, t-test for the onset of cursor movements and the onset of muscle activity, respectively), whereas the RT at the end of the compatible perturbation returned to baseline levels, as there was no significant difference to baseline (p = 0.923 and 0.917, t-test using cursor kinematics and muscle pattern for the RT determination, respectively). These results show that participants reacted significantly slower after an incompatible than after a compatible virtual surgery.

**Figure 9.**
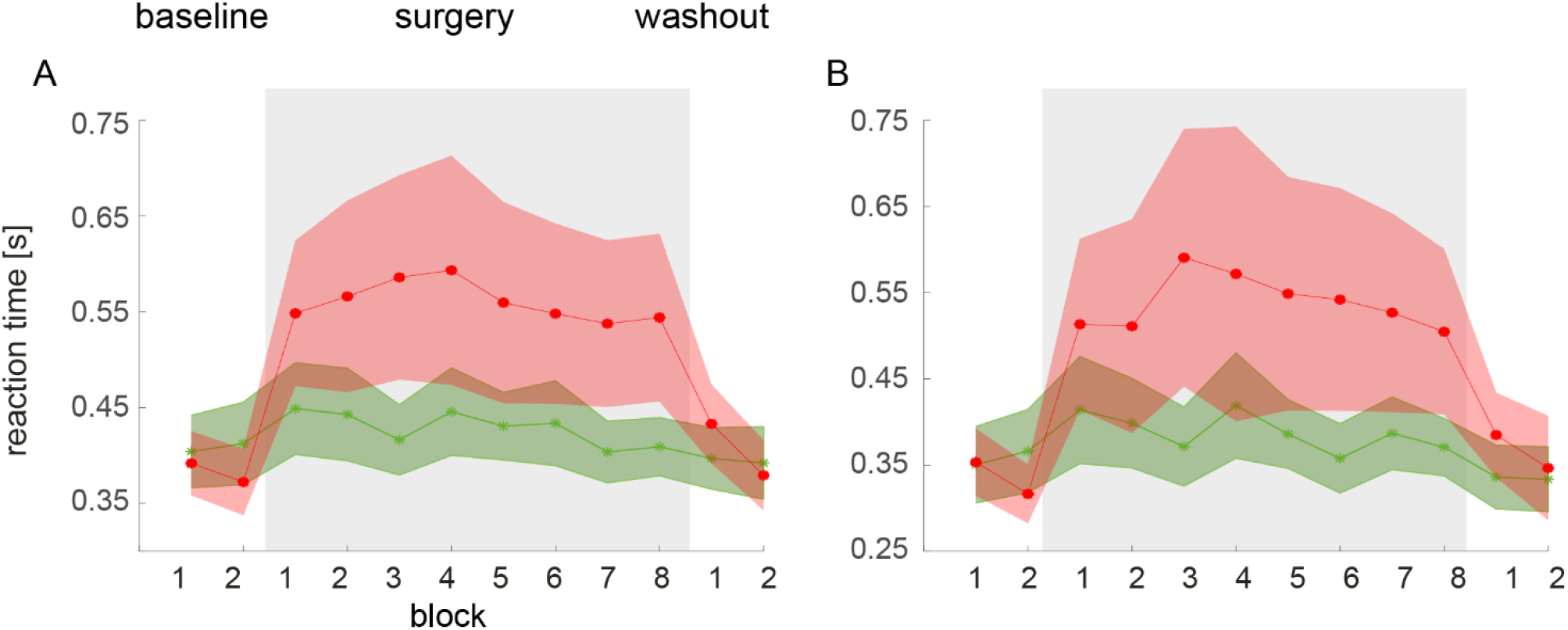
Comparison of reaction times after virtual surgeries. Reaction times using the cursor movement onset (A) and onset of muscle activity to determine the reaction times averaged across participants and blocks of 24 trials (solid lines, shaded areas indicate s.e.m.) for compatible virtual surgeries (*green*) and incompatible virtual surgeries (red).

## DISCUSSION

We investigated the acquisition of new muscle patterns after virtual surgeries, remapping of muscle forces using myoelectric control in a virtual environment. Healthy human participants improved the control of cursor movements in a reaching task after incompatible virtual surgeries in a single experimental session, thereby acquiring new muscle activity patterns. This result was made possible by allowing enough time for task-space exploration. Improvements in movement direction after incompatible surgeries occurred faster for corrective movements than for the initial movement, suggesting that activation of novel muscle patterns is more effective during feedback control, possibly due to an increased conscious awareness of the relevant online movement adjustments. Moreover, the significantly longer reaction time after incompatible virtual surgeries than after compatible virtual surgeries, further supports the use an explicit strategy for movement planning after an incompatible virtual surgery. Taken together, these results, on the one hand, indicate that exploration is an important factor for learning a control policy requiring the generation of novel muscle patterns and for motor skill learning and, on the other hand, suggest that human participants, with sufficient training, can learn new muscle synergies.

### Direct evidence for muscle synergies

Whether muscle synergies are the mechanism employed by the CNS to coordinate the activation of muscle groups to simplify motor control is debated (Kutch and Valero-Cuevas 2012; Tresch and Jarc 2009)\. One critical issue has been how to directly test the synergy hypothesis. Many studies have provided indirect support to the synergy hypothesis (d’Avella et al. 2006, 2008, 2011; d’Avella and Bizzi 2005; Cappellini et al. 2006; Chvatal and Ting 2012, 2013; D’Andola et al. 2013; De Marchis et al. 2018; Dominici et al. 2011; Frere and Hug 2012; Gentner et al. 2013; Hug et al. 2011; Ivanenko et al. 2007; Overduin et al. 2008; Roh et al. 2012; Torres-Oviedo and Ting 2007, 2010), as they only show that the observed muscle patterns lie in a low-dimensional space (d’Avella et al. 2003). Although low-dimensionality of the muscle patterns is the distinguishing feature of a modular controller, these studies do not provide a definitive answer on the neural origin of muscle synergies (Bizzi and Cheung 2013). The question remains of whether low-dimensionality is a mere consequence of how a non-modular controller solves the specific task or of the inherent biomechanical constraints (Kutch and Valero-Cuevas 2012; Steele et al. 2015), rather than the result of a neural organization of muscle synergies.

We have previously provided direct evidence for a neural origin of muscle synergies, by comparing the adaptation to perturbations incompatible with the muscle synergies and to perturbations that are compatible with them (Berger et al. 2013). By generating perturbations that could not be compensated by simply recombining pre-existing muscle synergies, we could investigate whether the participants were able to learn to use novel muscle activity patterns. Participants performed much worse during the incompatible than the compatible perturbations, a result predicted by the synergy hypothesis, under the assumption that learning novel muscle synergies is a slower process than adjusting the combination of existing synergies (Berger et al. 2013). However, a limitation of that study was that trial duration was short (2 seconds) and did not allow participants to explore the task space. In such conditions, participants reduced the initial direction error even after incompatible virtual surgeries, but they failed to successfully complete the task. We therefore hypothesized that, as with learning complex skills such as musical instrument playing, learning how to perform an isometric reaching task after an incompatible surgery would be possible with enough task space exploration. Thus, in this study, we investigated whether longer trial duration than in our previous study would affect the ability to overcome incompatible virtual surgeries. In the present study we extended our previous results showing, with a similar experimental protocol, that participants can learn to successfully complete the task even after incompatible surgeries, provided enough time for task exploration is available.

Recent studies have used the same conceptual approach that we have developed to probe the effect of manipulations that require activation of muscle patterns outside and within the manifold characterized by muscle synergies (d’Avella and Pai 2010; Berger et al. 2013) to assess neural constraints on learning. In studies using non-human primates (Oby et al. 2019; Sadtler et al. 2014), each neuron –rather than each muscle—is associated to a cursor movement direction, and perturbations affect the mapping between the neuronal activity and cursor movements. One important difference is that, while we mapped one of the dimensions in the muscle space effective in generating cursor movements (i.e., within the intrinsic manifold) completely into the null space (i.e., outside the intrinsic manifold), in the cited neural studies the remapped directions were not orthogonal to the intrinsic manifold. The authors stated that in this way they selected mappings that were difficult enough to require substantial learning but not too difficult such that the animals would give up. Despite these methodological differences, their results are in accordance with both our present and previous results (Berger et al., 2013).

### Effect of virtual surgeries on task performances

In our previous study, we had shown that participants performed much worse during incompatible than during compatible perturbations. Such result, supporting the synergy hypothesis, is reproduced in the present experiment, with longer trial durations. To prevent fatigue, the total number of trials in the current experiment was reduced by 33% (i.e., 192 vs. 288 trials per virtual surgery). Nonetheless, we still found a significant improvement in the initial direction error at the end of the exposure to the incompatible virtual surgery. The error in the initial direction of the cursor evaluates the open-loop performance, i.e., without any online, visually guided, feedback correction, reflecting the performance of a feedforward controller. Immediately after the initial movement, participants had up to 15 seconds in each trial to correct any inaccuracy and were allowed to use visual feedback and to gain experience of the relationship between muscle activation patterns and cursor movements. On average, participants were exposed to incompatible virtual surgeries for 25.1 min, and to compatible virtual surgeries for 16.6 min. After an incompatible surgery, participants had a larger initial path length in their trajectories, but the area spanned by the path was nevertheless smaller than after the compatible surgery. This observation is in accordance with the hypothesis that muscle patterns are generated by combining muscle synergies, as incompatible surgeries constrain the forces generated in the task space to one direction only (see Figure 2G).

In order to include also the effect of visual feedback on performance, we considered the fraction of trials in which the cursor did not reach and hold the target position in the available time. Similarly to the initial direction error, the fraction of unsuccessful trials revealed a better adaptation to the compatible virtual surgery than to the incompatible one, as predicted by the synergy hypothesis, and in line with our previous results (Berger et al. 2013). However, an important difference is that, in our previous study, at the end of the exposure to the incompatible surgery the relative number of successful trials did not show a significant increase. On the contrary, in the present study participants significantly increased the number of successful trials at the end of the incompatible perturbations. This result indicates that, despite a lower total number of trials, longer trial duration and visual feedback increase the participants’ ability to adapt to an incompatible perturbation. The fact that participants significantly improved their trial success rate only when more time was available and visual feedback could be used for online corrections suggests that exploration is an important factor for learning a new control policy overcoming the perturbation.

Peaks in the speed profile are commonly used as indicators of movement corrections. In our data, the numbers of speed peaks increased significantly after both types of surgeries. This reflects the fact that participants made use of visual feedback to correct online the movement of the cursor. Furthermore, the number of corrective movements decreased significantly during the exposure to both compatible and incompatible virtual surgeries. Multiple online corrections, together with the significant improvement of trial success, suggests that participants exploited visual feedback to overcome the incompatible perturbation. By the way, at the end of the exposure to both surgery types the number of speed peaks was significantly higher than before exposure, reflecting the fact that also after the compatible virtual surgery online corrections had not completely adapted to baseline levels.

#### Feedback and feedforward control processes after incompatible and compatible virtual surgeries

Our results on the direction error of the initial movement and first corrective movement helped us to gain new insights into the adaptation of both feedforward (i.e., offline adaptation, obtained by adjusting the cursor movement on the following trial) and feedback control (i.e., online control, obtained through cursor movement adjustments during execution) after the two types of virtual surgeries. During both perturbations, participants adapted their movements and improved their reaching performance. However, we found remarkable differences in the time course of adaptation of feedforward and feedback control processes between the two types of perturbations. Feedforward control –assessed by the initial direction error– improved after both virtual surgeries over time, and it did so significantly faster after the exposure to a compatible virtual surgery than to an incompatible one, in accordance with the synergy hypotheses (as discussed above), and reproducing our previous results in Berger et al. (2013). Feedback control –assessed by the direction error of the first movement correction– also improved after both types of virtual surgeries. However, the initial direction error and the direction error of the corrective movement followed different time courses in the two types of surgeries. During the exposure to a compatible virtual surgery, subjects were able, immediately after the introduction of the surgery, to efficiently use feedback control. This led to an immediate, significant improvement of direction error of the first corrective movement. This result is in line with the observation that feedback control during learning of visuomotor rotations does not require adaptation (Kasuga et al. 2015; Telgen et al. 2014). Rather, it has been suggested that visuomotor mapping is adapted by adding some part of the corrective response under the old mapping to the known motor command (Kawato and Gomi 1992). Towards the end of the exposure to a compatible virtual surgery, the initial direction error was similar to the error after the first movement correction, showing that participants improved significantly the effectiveness of feedforward control throughout the surgery with similar performance as when using feedback control. On the contrary, immediately after incompatible virtual surgeries, subjects were unable to efficiently adjust the cursor movements, and online corrections were inaccurate. This observation strongly contrasts to the observation of a moderate direction error for online corrections after compatible virtual surgeries. Reaching towards a target after an incompatible virtual surgery is challenging, as no muscle activation pattern generated as combination of the existing synergies (i.e., in the synergy space) can lead to an improvement in performance. During the time course of the exposure to the virtual surgery the direction error of both the initial and the first corrective movement improved. This points to subjects being able to activate muscle patterns outside the synergy space, which can also be seen in the results on muscle reconstruction (Figures 5 and 6). The fact that direction error improved significantly faster for the first movement correction than for the initial movement, suggests that participants learned to use more efficiently feedback mechanisms with increasing exposure to the incompatible perturbation, suggesting that activation of novel muscle patterns is more effective during online guided feedback control, possibly due to some underlying strategic process being employed due to an increased conscious awareness of the relevant online movement adjustments. Participants also significantly improved their initial movement directions, showing also a more accurate movement planning. Taken together, these results show that participants adapted to the incompatible virtual surgery by adjusting both feedforward and feedback control mechanisms.

Little is known how feedforward and feedback control relate to each other and its relationship is still a matter of debate. A number of studies have shown that feedback and feedforward control processes are tightly coupled (Ahmadi-Pajouh et al. 2012; Cluff and Scott 2013; Wagner and Smith 2008). In contrast, our results show a dissociation between movement planning and online correction mechanisms after incompatible virtual surgeries with different time courses of the adaptation of feedforward and feedback control processes compared to those observed after compatible virtual surgeries. The two processes are at least partially independent from each other, as they have different time courses. However, our results are in line with several studies using a mirror-reversed visual environment (Gritsenko and Kalaska 2010; Kasuga et al. 2015; Kuang and Gail 2015). Gritsenko & Kalaska (2010) showed that subjects can selectively suppress rapid online correction under reversed vision, suggesting that when exposed to reversed visual input, subjects do not adapt their motor planning but rather their motor control (Kuang and Gail 2015). Kasuga et al. (2015) showed that under reversed vision, feedback control and forward control are learned separately, as feedback control only improved when continuous visual feedback was provided and online movement corrections were allowed. In contrast, feedforward control improved more when online corrections were not allowed. Thus, the seemingly contrasting results supporting coupled versus independent feedforward and feedback control processes might depend on whether the task requires learning of a control policy de novo, e.g., when learning a new skill, in which case feedforward and feedback mechanisms might be learnt independently (Kasuga et al. 2015; Kawato and Gomi 1992), or whether an existing control policy is adapted, as in the case of visuomotor rotation. With this line of reasoning, the differences in feedforward and feedback control adaptation between the two types of virtual surgeries might be explained by the coupling of the two processes during motor adaptation, as after compatible virtual surgeries only the synergy coefficients need to be adjusted, and their partial uncoupling after incompatible virtual surgeries, when a new control policy needs to be acquired.

Subjects were better at compensating an incompatible perturbation when generating online corrections than at the beginning of the subsequent trial, even at the end of the surgery. Why subjects were able to improve the direction error of the cursor better during feedback then during feedforward control after incompatible virtual surgery is not completely clear. However, this observation suggests that the exploration of new muscle patterns required for overcoming an incompatible virtual surgery may be more easily achieved during online control. A possible explanation might be that subjects quickly become aware of the need to explore through online movement adjustments. Thus, a contribution of an explicit strategy may be an important factor for learning a new control policy required to overcome an incompatible virtual surgery.

#### Implicit and explicit control strategies after incompatible and compatible virtual surgeries

It is commonly believed that reaction time (RT) reflects the underlying computations required for preparing an action, i.e., the time needed to complete the perceptual and motor-planning computations required to prepare an appropriate movement (Donders 1969; Friston et al. 1996; Sternberg 1969). Additional time for movement initiation is required when a new motor command is needed to achieve a given goal, such as during visuomotor adaptation that is reflected by an increased RT (Fernandez-Ruiz et al. 2011; Saijo and Gomi 2010). The increase in RT that we observed after compatible virtual surgeries (Figure 9) is in line with what has been reported after visuomotor rotations, as adaptation to visuomotor rotations can be seen as a type of compatible virtual surgery, since both only require adjusting the synergy coefficients while using the existing subject-specific muscle synergies. Remarkably, after incompatible virtual surgeries participants reacted significantly slower than after compatible virtual surgeries.

An important idea in motor skill learning research is that motor learning proceeds from predominantly explicit to implicit strategies as a learner develops from novice to expert (Anderson 1982; Fitts and Posner 1967; Miyamoto et al. 2020). An increased cognitive load due to the usage of an explicit strategy has been shown to have a stronger impact on RT than during implicit adaptation (Benson et al. 2011a). Moreover, Telgen and collaborators (Telgen et al. 2014), suggested that when the motor system acquires a new control policy, extra processing time is required. As incompatible virtual surgeries require the establishment of a new control policy, this should then consequently entail initially slower and possibly more explicit components (Hikosaka et al. 2002; Telgen et al. 2014), requiring longer processing time than for recalibrating a well-learned control policy. Thus, longer RTs after incompatible with respect to compatible virtual surgeries may be due to the fact that subjects relied more on an explicit strategy, such as cognitive reinterpretations of sensory signal and an increased muscle space exploration, possibly involving increased amount of voluntary co-contraction (Borzelli et al., 2018).

The hallmark of implicit learning during motor adaptation is the presence of after-effects. Persistence of motor changes, even when the perturbation is removed, is thought to reflect a change in the internal model (Wolpert et al. 1995). Consequently, the presence of an explicit, cognitive strategy may explain the absence of significant after-effects (Benson et al. 2011; Buch et al. 2003). Thus, further support to the hypothesis that subjects relied on an explicit learning strategy to overcome the incompatible virtual surgery is given by the reduced after-effects after incompatible virtual surgeries (Figure 4A) compared to after compatible virtual surgeries. Taken together, our results provide evidence for different adaptation processes involved in overcoming an incompatible or a compatible virtual surgery. They suggest that incompatible virtual surgeries require an explicit exploratory mechanism for synergy learning, leading to higher RTs and smaller after-effects with respect to compatible virtual surgeries, for which only combination coefficients of existing synergies need to be adjusted.

### Distinct learning processes for compatible and incompatible virtual surgeries

During a motor adaptation task such as visuomotor rotation learning, online motor control relies upon our predictions about the consequences of our movement commands as well as upon feedback received during movement execution. When a movement misses its target, the brain quickly adapts the next motor command to prevent further errors. Computational models of such motor adaptation tasks presume a previously existing control policy that is then recalibrated to minimize the sensory-prediction errors (Diedrichsen 2007; Wolpert and Flanagan 2016). After compatible virtual surgeries (like in visuomotor rotations as they can be seen as a type of compatible perturbation), cursor trajectories that can be generated within the know repertoire of motor commands -- by recombining existing muscle synergies -- provide information about the novel mapping between motor commands and task space and the inaccurate cursor trajectories can thereby be used to improve performance in the following trials by calculating the error between the estimated trajectory and the actual trajectory. More specifically, in a synergy-based controller (Hagio and Kouzaki 2018), a compatible virtual surgery can be overcome by using existing muscle synergies and associating new synergy combinations to the movement targets. The hypothesis is that a new mapping of visual targets to synergy coefficients is learned and that such adaptive process occurs faster than the learning of new synergies. This would explain improved planning and offline corrections even in the experiments with shorter trial durations. This hypothesis is supported by the fact that participants were immediately able to use efficient feedback control to decrease the angular error. Interestingly, however, after compatible virtual surgeries, independent of trial duration, participants did not adapt completely and they failed to return to baseline, as the results of the initial direction error and the trial success rate demonstrate.

In contrast, during incompatible perturbations, as no combination of the existing muscle synergies can be used to move the cursor in all directions, participants are required to acquire a novel control policy, mapping targets onto new muscle patterns, to gain control over the entire task space. This means that the muscle combinations required to move the cursor in all directions are not in the repertoire of know actions, i.e., cannot be generated by recombination of existing muscle synergies. Thus, cursor trajectories that rely on the existing muscle synergies do not provide information about the novel mapping between motor commands and task space. Novel muscle patterns must be generated and both new synergies and a new mapping of visual targets into combination coefficients of these synergies need to be learned to overcome the perturbation. Here trial duration plays a key role. With a short trial duration, most participants are limited in their movements and are unable to successfully guide the cursor into the target. In contrast, with a longer trial duration, participants can successfully guide the cursor into the target. We thus argue that the inability of overcoming the incompatible perturbation in the previous experiment was due to shorter trial duration and consequent lack of exploration. Having more time to explore the task space thanks to longer trial duration, as in the present experiments, leads to gathering of more information about the effect of novel mapping in the task space which drives the adaptive processes required to overcome the virtual surgeries. This hypothesis is supported by our results of the movement direction during incompatible perturbations, which showed that subjects rely more heavily on movement corrections to improve the cursor movement direction, possibly due to an increased conscious awareness of the relevant online movement adjustments. Thus, in addition to learning of new mappings of visual targets into synergy coefficients occurring after both compatible and incompatible surgeries, learning of new muscle synergies is required to adapt to an incompatible virtual surgery. This second process may be responsible for changing the structure of the synergies to recover their capability of generating forces in all directions. The establishment of a new control policy should entail initially slower components (Hikosaka et al. 2002) and requiring additional processing time (Telgen et al. 2014). The exploration of new muscle coordination patterns and the acquisition of new task-specific synergies required by incompatible virtual surgeries may be realized through goal babbling (Rolf et al. 2010), which main idea is to generate knowledge by exploration or by goal-related feedback mechanism (Rohde et al. 2019). We suggest that such a process is engaged only when the existing synergies are unable to perform a task, such as after a major change in the musculoskeletal system due to injury or when learning a new motor skill. Our present results provide supporting evidence for this hypothesis (see the change in EMG patterns during the incompatible perturbation of Figure 6). Indeed, EMG patterns significantly changed during the incompatible perturbations, while after the compatible virtual surgery the similarity of the reconstruction remained high throughout the whole perturbation period. In this case, existing muscle activity patterns were simply re-associated with new movements.

### Conclusions

In summary, we showed that adaptation to virtual surgeries incompatible to the subject-specific synergies is possible, provided sufficient time for task space exploration is given. Novel combinations of muscle activity patterns, potentially previously unknown to the CNS, can be put synergistically together. Our results suggest that learning new synergies is related to the exceptional human capacity for acquiring a wide variety of novel motor skills with practice.

The mechanisms of the neural implementation of muscle synergies require additional experiments with patients and healthy participants, possibly combined with stimulation. An interesting avenue for future research would be to determine if the constraints we observed on the muscle patterns remain over even longer timescales. As with musical instrument playing, learning a motor skill and the underlying muscle synergies might require several days, or months, of practice.

## ACKNOWLEDGMENTS

We thank F. Lacquaniti and A. Pazienti, for helpful comments on an earlier version of the manuscript. This work was supported by the Human Frontier Science Program Organization (RGP11/2008), the European Community’s Seventh Framework Programme (FP7/2007–2013-Challenge 2-Cognitive Systems, Interaction, Robotics, grant agreement No 248311-AMARSi), Italian Space Agency (ASI-MARS-PRE DC-VUM-2017-006), Italian University Ministry (PRIN grants 2015HFWRYY_001 and 2017CBF8NJ_005), and the Italian Ministry of Health Grant RF-2019-12370271.

## Grants

This work was supported by the Human Frontier Science Program Organization (RGP11/2008), the European Community’s Seventh Framework Programme (FP7/2007–2013-Challenge 2-Cognitive Systems, Interaction, Robotics, grant agreement No 248311-AMARSi), Italian Space Agency (ASI-MARS-PRE DC-VUM-2017-006), Italian University Ministry (PRIN grants 2015HFWRYY_001 and 2017CBF8NJ_005), and the Italian Ministry of Health Grant RF-2019-12370271.

## Disclosures

The authors declare no competing financial interests.

## Notes

### Competing Interest Statement

The authors have declared no competing interest.

